# Multi-omics approach identifies germline regulatory variants associated with hematopoietic malignancies in retriever dog breeds

**DOI:** 10.1101/2021.04.05.438235

**Authors:** Jacquelyn M. Evans, Heidi G. Parker, Gerard R. Rutteman, Jocelyn Plassais, Guy CM Grinwis, Alexander C. Harris, Susan E. Lana, Elaine A. Ostrander

**Author notes:** Corresponding Author (EAO).

## Abstract

Histiocytic sarcoma is an aggressive hematopoietic malignancy of mature tissue histiocytes with a poorly understood etiology in humans. A histologically and clinically similar counterpart affects flat-coated retrievers (FCRs) at unusually high frequency, with 20% developing the lethal disease. The similar clinical presentation combined with the closed population structure of dogs, leading to high genetic homogeneity, makes dogs an excellent model for genetic studies of cancer susceptibility. To determine the genetic risk factors underlying histiocytic sarcoma in FCRs, we conducted multiple genome-wide association studies (GWASs), identifying two loci that confer significant risk on canine chromosomes (CFA) 5 (*P*_wald_=4.83×10^−9^) and 19 (*P*_wald_=2.25×10^−7^). We subsequently undertook a multi-omics approach that has been largely unexplored in the canine model to interrogate these regions, generating whole genome, transcriptome, and chromatin immunoprecipitation sequencing. These data highlight the PI3K pathway gene *PIK3R6* on CFA5, and proximal candidate regulatory variants that are strongly associated with histiocytic sarcoma and predicted to impact transcription factor binding. The CFA5 association colocalizes with susceptibility loci for two hematopoietic malignancies, hemangiosarcoma and B-cell lymphoma, in the closely related golden retriever breed, revealing the risk contribution this single locus makes to multiple hematological cancers. By comparison, the CFA19 locus is unique to the FCR and harbors risk alleles associated with upregulation of *TNFAIP6*, which itself affects cell migration and metastasis. Together, these loci explain ~35% of disease risk, an exceptionally high value that demonstrates the advantages of domestic dogs for complex trait mapping and genetic studies of cancer susceptibility.

## Introduction

Histiocytic sarcoma is a rare, aggressive cancer of dendritic cells and macrophages that accounts for < 1% of hematopoietic malignancies in humans (1, 2). Tumors arise as the primary neoplasm or concurrently with other hematological malignancies, such as lymphoma or chronic lymphocytic leukemia, through transdifferentiation or a common neoplastic precursor (3). The disease is diagnosed most frequently in adulthood and may present as localized or disseminated, with tumors in multiple sites, including the spleen, liver, lymph nodes, gastrointestinal tract, and skin (2). The neoplastic cells are typically large and round-polygonal in shape; spindle cells may also be present (2). Treatment response is poor, and most patients succumb to the disease within two years (2). Limited biological samples for this rare cancer have hindered large-scale genetic studies.

A histologically and clinically similar disease, also termed histiocytic sarcoma, occurs spontaneously in dogs (4, 5). Although rare across breeds as a whole, histiocytic sarcoma is common in flat-coated retrievers (FCRs) and Bernese mountain dogs, affecting ~20% and 25% of dogs in each breed with near uniform fatality (6-8). The canine disease also presents as localized in periarticular tissue or disseminated in the viscera, with the former more common in the FCR (9) and the latter typical in the Bernese mountain dog (7).

Dogs are a well-described, naturally-occurring model for many human cancers including sarcomas and hematological cancers, such as osteosarcoma, lymphoma, and leukemia (10-12). Most breeds were developed within the last 200 years (13), and population bottlenecks coupled with strong selection for morphological and behavioral traits has created a unique population structure characterized by reduced genetic diversity and long haplotype blocks within breeds, but also substantial across-breed variation (14, 15). Detrimental alleles have become enriched as a consequence of breed formation processes, leading to disease predisposition. These characteristics facilitate complex trait mapping studies in dogs, which require roughly two orders of magnitude fewer markers compared to human GWASs and as little as 200 individuals (13-15).

The FCR and Bernese mountain dog are ideal for gaining insight into the etiology of histiocytic sarcoma, as the unusually strong breed prevalence suggests a highly penetrant heritable component. While the two breeds do not share recent common ancestors and they were bred for distinct physical attributes (16), the similarity of disease progression and outcomes, as histiocytic sarcoma always progresses to metastatic disease with full lethality, is the same between the breeds. The overall rarity of the disease, and its total absence in the majority of domestic dog breeds, argues for at least partially overlapping genetic risk factors, likely reflecting the explosion of modern breeds in Western Europe <200 years ago. However, as the primary differences between affected dogs appear to be in initial disease presentation, it is likely that at least partially independent genetic mechanisms underly histiocytic sarcoma susceptibility in each. While our previous GWAS in Bernese mountain dogs successfully identified a locus on canine chromosome (CFA) 11 (17), the FCR genetic predisposition has been hitherto unexplored.

Here, we conducted multiple GWASs in FCRs, identifying two loci on CFA5 and 19 that confer risk for histiocytic sarcoma. We advance beyond previous cancer and complex disease studies in the dog, analyzing whole genome, transcriptome, and ChIP sequencing data to identify putative regulatory variants associated with histiocytic sarcoma susceptibility in FCRs that may also confer risk for hematopoietic cancers in other breeds. We thus leverage the advantages of the canine model system to further our understanding of the biology of a rare human cancer with implications for both canine and human health.

## Results

### CFA5 and CFA19 confer risk for histiocytic sarcoma

A GWAS including 177 FCR histiocytic sarcoma cases and 132 FCR controls (Table 1) was performed using 108 084 SNPs. Principal component analysis revealed stratification between FCRs of European vs. North American origin (Fig 1). GWAS was performed in GEMMA (18) using a kinship matrix and linear mixed model to correct for population structure with a genomic inflation factor (λ) of 0.97. A single association exceeding Bonferroni significance (4.26×10^−7^) was identified on CFA5 with *P*_wald_=4.83×10^−9^ (Fig 2A, Supplementary Table S1). The top 27 markers are in high linkage disequilibrium (LD; r2≥0.8) and span a 4.3 Mb region around the lead SNP (CFA5:33001550) (Fig 2B). A shared haplotype was identified among 90% of cases, with recombination events defining a narrower 1.2 Mb FCR risk haplotype at CFA5:32389061-33633274 (Fig 2C; Supplementary Table S2), which harbors over 40 genes. The risk haplotype is also present in 64% of control dogs. There is considerable LD among markers at this locus (r^2^ ≥ 0.6 28-37 Mb) with the broader GWAS signal extending to 28 Mb, and many cases continue to share a common haplotype throughout the region.

**Table 1.** Description of samples used in each analysis.

| **Analysis** | **Samples** |
| --- | --- |
| FCR histiocytic sarcoma GWAS #1 | 177 FCR cases, 132 FCR controls |
| FCR histiocytic sarcoma GWAS #2 - heterozygotes for CFA5 risk | 94 FCR cases, 43 FCR controls (subset of GWAS #1) |
| RNA-seq analyses | 11 FCR blood samples |
| H3K4me1 and H3K4me3 ChIP-seq | 7 canine blood samples |
| Whole genome sequence filtering | 3 FCR histiocytic sarcoma cases, 1 FCR control, 4 golden retriever B-cell lymphoma cases<sup>a</sup> |
| Candidate variant genotyping - histiocytic sarcoma association | Minimum of 79 cases and 69 controls from the GWAS cohort |
| Variant genotyping - retrievers with hematopoietic cancer | FCR lymphoma cases (n=20); golden retriever B-cell lymphoma (n=9) and histiocytic sarcoma (n=21) cases |

<sup>a</sup>Publicly-available genomes from (19)

**Fig 1.**
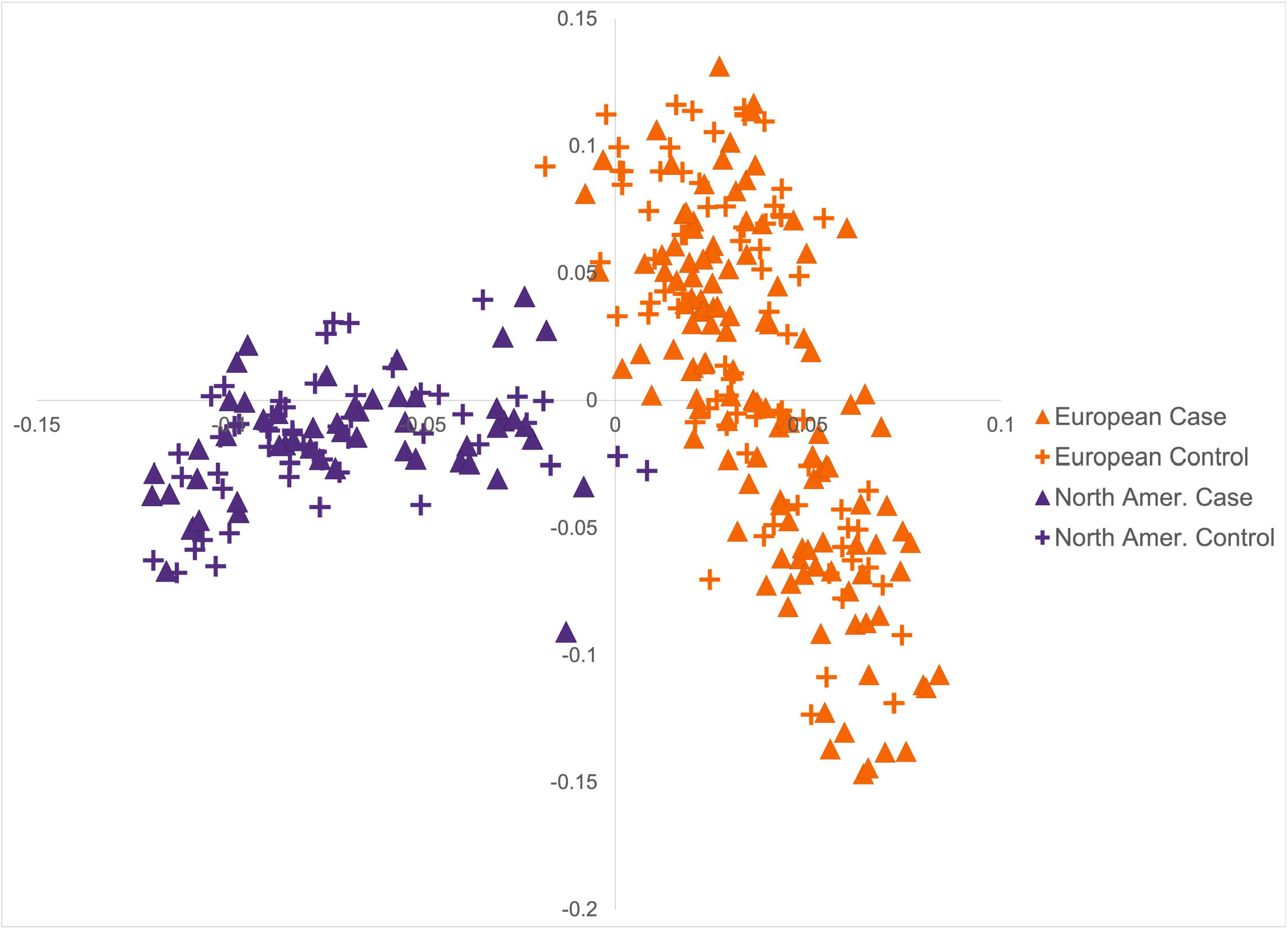
Principal components analysis of FCR GWAS. Principal components 1 (13.6% variance) and 2 (6.6% variance) are plotted on the x and y-axes, respectively. The European and North American FCRs (n=309) form subpopulations with cases and controls distributed throughout both groups.

**Fig 2.**
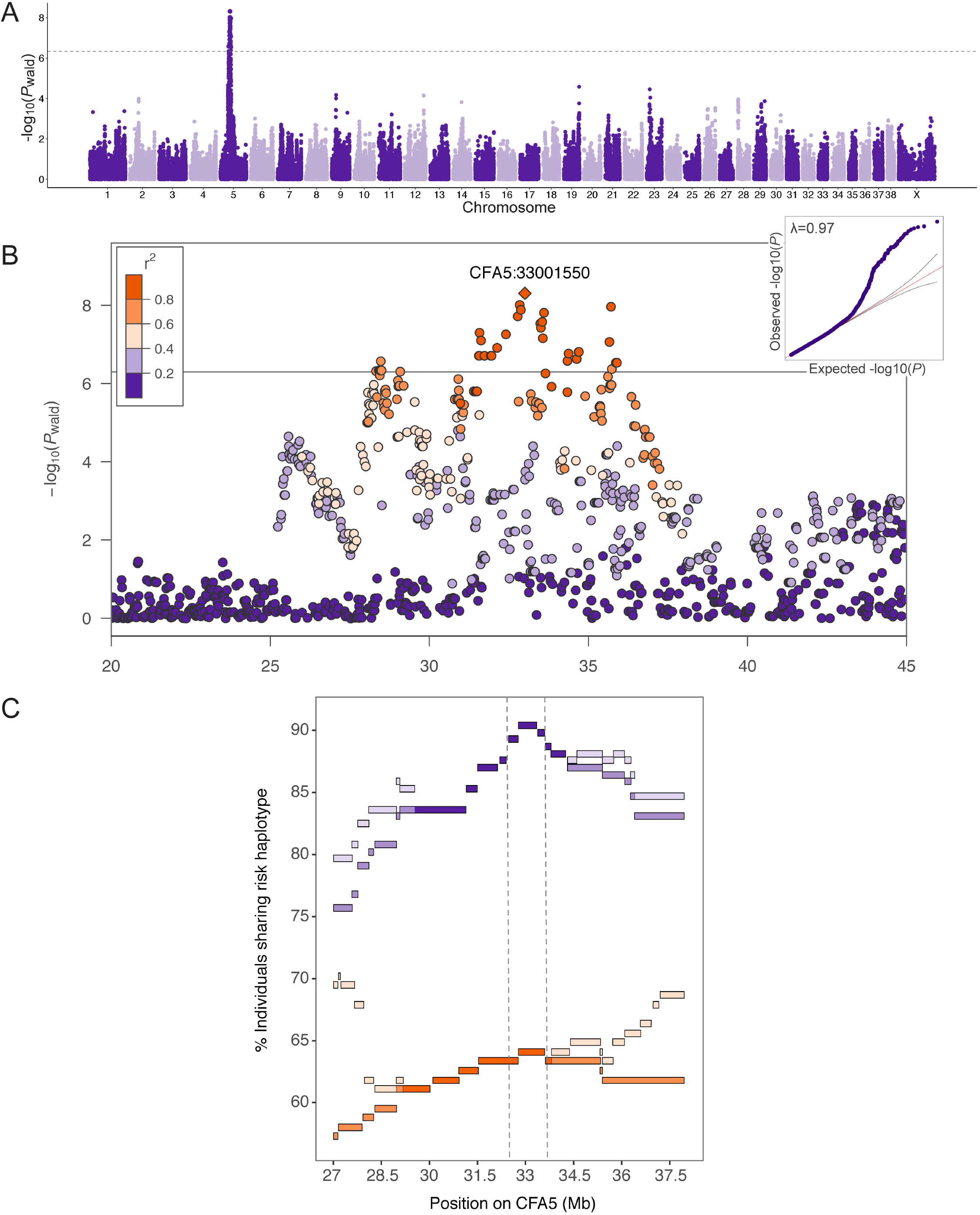
Genome-wide association study results for 177 histiocytic sarcoma FCR cases and 132 controls. A) Manhattan plot of -log_10_*P*-values (y-axis) for 108084 Illumina SNPs plotted against chromosome position in CanFam3.1 (x-axis). The Bonferroni threshold is plotted as a solid line (-log_10_*P*=6.33). B) Regional Manhattan plot of the CFA5 association with SNPs color-coded according to pairwise LD (r^2^) with the lead SNP. C) Length of risk haplotype sharing among cases (purple) and controls (orange) is plotted on the x-axis with the percentage of dogs sharing on the y-axis. Continuous loss of haplotype sharing is tracked in darker purple/orange, while the lighter shades mark points at which some individuals re-gain the common risk haplotype.

The CFA5 risk haplotype is present in the heterozygous state in 53% of cases and 43% of controls. To determine whether additional loci differentiate these groups, we performed a GWAS using only cases and controls heterozygous for the CFA5 risk haplotype (94 vs. 43; Fig 3, Table 1), thereby neutralizing the effect of the CFA5 locus. To reduce the possibility that our control group contained dogs who could eventually develop histiocytic sarcoma, we applied a more stringent minimum age at collection for controls (11 years), discarding samples from dogs in the lowest age quartile. This provided further separation between controls and cases, as 75% of cases were diagnosed at <10 years of age, while preserving as much power as possible for the GWAS (Supplementary Fig S1, Supplementary Table S3). A single locus at 52 Mb on CFA19 (*P*_wald_=2.25×10^−7^) exceeded Bonferroni significance (4.67×10^−7^) and was confirmed after permutations (*P*_permutations_=0.014). This approach produced a more robust association compared to that which includes CFA5 genotypes as a covariate in the total GWAS cohort (CFA19:52487724 *P*_wald_=4.25×10^−5^, Supplementary Table S4). A 741 kb critical interval is demarcated by the flanking SNPs in highest LD with the lead SNP (CFA19:52487724, r^2^≥0.6), encompassing just three genes. Ninety-nine of the 177 cases in the total GWAS cohort (n=309) had periarticular tumors and 77 had tumors in other locations at the time of diagnosis. The risk allele at the CFA19 locus was more common among periarticular cases (*P*_Fisher_=0.015, OR=2.78, 95%CI=1.21-6.37).

**Fig 3.**
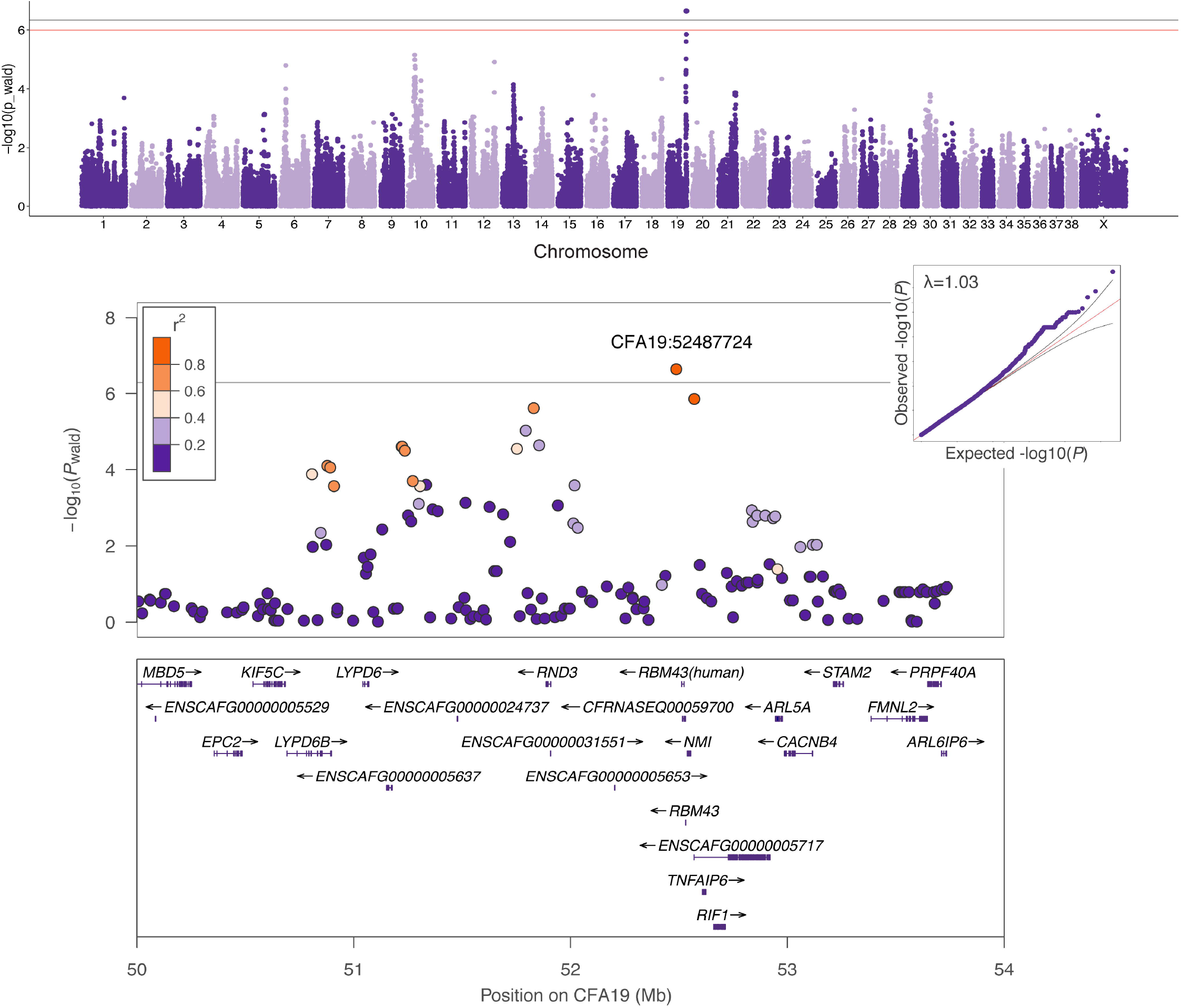
Genome-wide association study results for 94 histiocytic sarcoma FCR cases vs. 43 controls. All cases and controls were heterozygous for the CFA5 risk haplotype. Manhattan plot of -log_10_*P*-values (y-axis) for 107102 SNPs by chromosome position in CanFam3.1 (x-axis) is shown at the top. The Bonferroni and 5% permutations thresholds are plotted as gray and red lines, respectively. QQ plot with genomic inflation factor (λ) and regional Manhattan plot of CFA19 locus, showing pairwise LD (r^2^) relative to the lead SNP are below with genes in the region plotted at the bottom.

In the total cohort (n=309), SNP genotypes from all autosomes explain 27% ± 14% *P*_LRT_=0.0034) of the risk for developing histiocytic sarcoma. The CFA5:25-40Mb locus alone explains 22% ± 13% *P*=1.15×10^−5^, while CFA19:50.5-53Mb explains 8% ± 5% *P*_LRT_=2.05×10^−4^. Together, these loci account for 35-37% ± 13% of the phenotypic variance (*P*_LRT_=1.44×10^−8^). When considering CFA5 and CFA19 genotypes in combination, 39% of dogs who are heterozygous at both loci are cases, whereas 80% of dogs heterozygous at CFA5 and homozygous at CFA19 are cases (Table 2). Thus, when CFA19 data are included, we observe greatly improved separation of cases and controls relative to analysis with CFA5 genotypes alone.

**Table 2.** Genotypic combinations for CFA5 and CFA19 risk loci in FCR cases and controls.

| Genotype<sup>a</sup> | Cases (n=177) | Controls (n=98)<sup>b</sup> | Penetrance (%)<sup>c</sup> |
| --- | --- | --- | --- |
| aabb | 0 | 1 | - |
| aaBb | 3 | 7 | 30 |
| aaBB<sup>d</sup> | 14 | 29 | 33 |
| Aabb | 0 | 5 | - |
| AaBb | 11 | 17 | 39 |
| AaBB<sup>d</sup> | 83 | 21 | 80 |
| AAbb | 1 | 0 | - |
| AABb | 11 | 6 | 65 |
| AABB<sup>d</sup> | 54 | 12 | 82 |

<sup>a</sup>“A” represents the risk haplotype at CFA5; “a” is any non-risk haplotype. “B” represents the risk allele at the CFA19 lead SNP; “b” is non-risk.
<sup>b</sup>Controls were $\geq 11$ years of age at sample collection.
<sup>c</sup>The proportion of dogs with each genotype who are cases.
<sup>d</sup>These genotype combinations were present in FCRs used for WGS variant filtration.

### Multiple hematopoietic malignancies are associated with CFA5 locus

The CFA5 region colocalizes with previously-identified associations for two common hematological malignancies in golden retrievers: hemangiosarcoma (29 Mb) and B-cell lymphoma (34 Mb; Fig 4A) (19). Although distinct cancers, histiocytic sarcoma, B-cell lymphoma, and hemangiosarcoma all arise from cells in the hematopoietic stem cell pathway: dendritic cells and macrophages, B lymphocytes, and hematopoietic precursor cells, respectively (4, 8, 20, 21). FCRs and golden retrievers are closely related breeds, sharing an immediate common ancestor among the retriever phylogenetic clade (16). To search for shared risk haplotypes at this locus, we examined published genotypes (19) from golden retrievers diagnosed with hemangiosarcoma or B-cell lymphoma. Using the same haplotype analysis applied to FCRs (See Methods), we defined a 1.4 Mb B-cell lymphoma risk haplotype (CFA5:33001663-34362236) encompassing the lead golden retriever GWAS SNP (CFA5:34117726) for this cancer (Supplementary Table S2). This haplotype overlaps the FCR risk haplotype for 631 kb (Fig 4B), and the interval is strongly associated with hematopoietic cancer in both breeds, with a combined *P*-value of 4.17×10^−10^ compared to 3.43×10^−7^ and 2.00×10^−4^ in FCRs and golden retrievers alone, respectively. Direct overlap between golden retriever and FCR haplotypes was not observed at the 29 Mb hemangiosarcoma risk locus.

**Fig 4.**
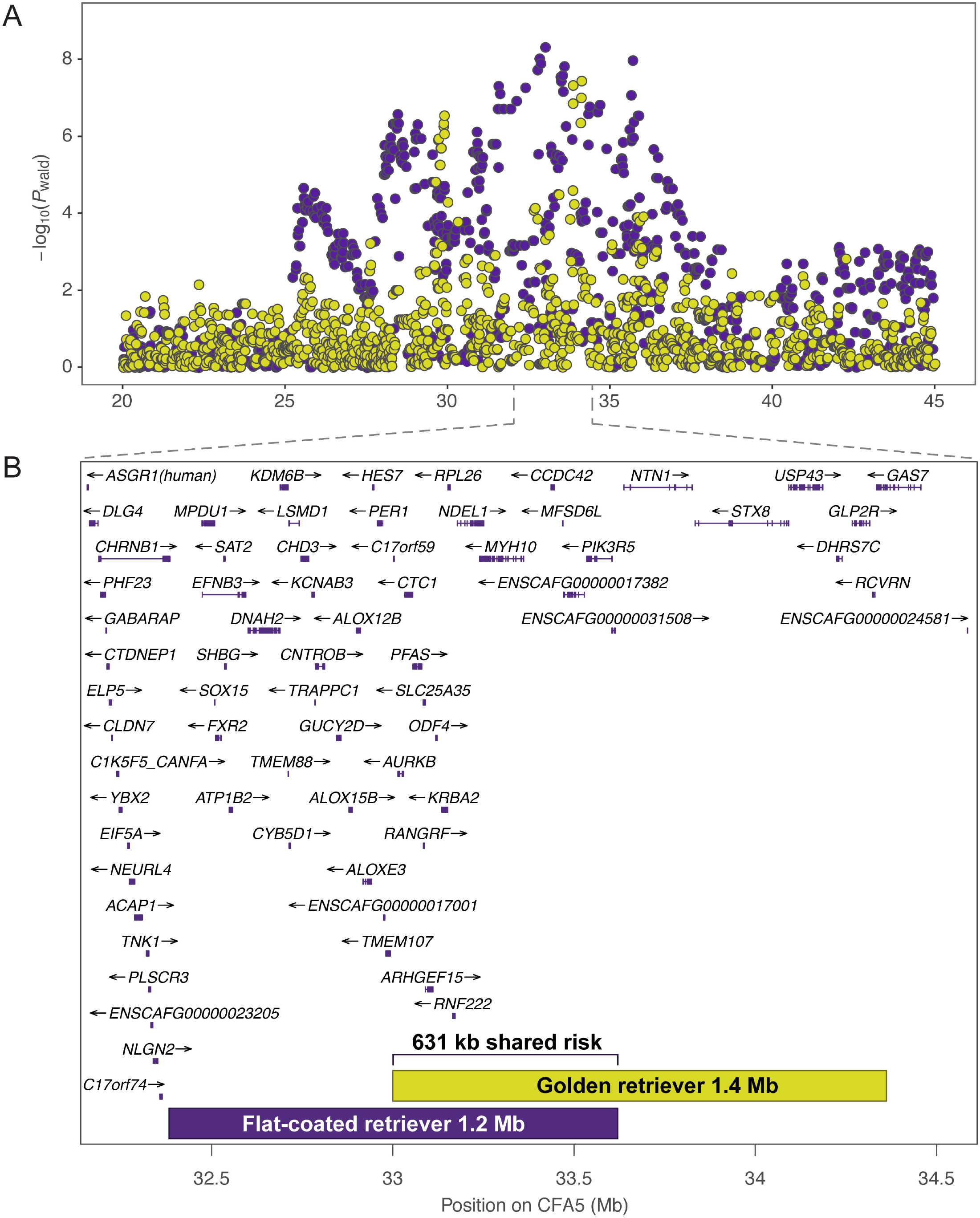
Colocalization of FCR histiocytic sarcoma CFA5 locus with golden retriever hematological malignancy loci. A) Regional Manhattan plot showing FCR histiocytic sarcoma GWAS SNPs in purple. Results of a combined golden retriever GWAS with 142 hemangiosarcoma, 41 B-cell lymphoma, and 172 controls are overlaid in gold. The peaks at 29 Mb and 33 Mb (CanFam3.1) in golden retrievers correspond to hemangiosarcoma and B-cell lymphoma risk, respectively. B) Regions harboring risk haplotypes identified independently in FCRs (purple) with histiocytic sarcoma and golden retrievers (gold) with B-cell lymphoma are plotted with genes in the region. The risk haplotypes overlap for a shared 631 kb span (bracket).

### RNA-seq and allele-specific expression

To investigate potential effects of the CFA5 risk haplotype on gene expression, RNA-seq data were generated from RNA isolated from 11 FCR whole blood samples (Table 1). Differential expression analysis was based on the risk haplotype, comparing four dogs who were homozygous for the risk haplotype vs. seven dogs who were heterozygous. The frequency of the risk allele in the FCR control population indicates the difficulty in finding homozygous non-risk individuals; however, this would clearly be beneficial in future expression studies. Forty-three genes and five non-coding RNAs demonstrated significant differential expression. The nearest gene to the CFA5 critical interval, *NLRP1*, was 1.7Mb upstream, suggesting the risk locus may have distal effects (Supplementary Table S5). When comparing gene expression levels in individual samples to the average expression across controls (see Methods, (22)), seven genes and one lncRNA demonstrated significant individual expression (z-score≥|2.5|) among cases. After excluding one heterozygous dog who received chemotherapy one week prior to the blood draw, comparison of the four dogs homozygous for the CFA5 risk haplotype to the remaining six heterozygous dogs revealed an additional 17 genes or non-coding RNAs with significant differential and individual expression (Supplementary Table S5).

Because RNA samples were only available from a small number of homozygous and heterozygous individuals and no dogs without risk alleles, the power to detect changes in gene regulation through differential expression analyses was limited. Allele-specific expression (ASE) analysis provides an alternative approach to investigate differential expression utilizing heterozygous individuals. ASE compares expression levels for two alleles at a given coding SNP within an individual, which may result from *cis*-regulation by variants in non-coding regions. This controls for sources of error between individuals, like environmental, technical, or *trans*-regulatory effects (23, 24). We performed an ASE analysis for RNA samples isolated from blood for the seven FCRs heterozygous for the CFA5 risk haplotype. We examined genes within 500 kb on either side of the 631 kb shared risk haplotype, extending our search to include potential long-range enhancer-gene interactions (25). Variants demonstrating significant ASE in two or more FCRs were identified in seven genes: *CD68, MPDU1, CHD3, BORCS6, NDEL1*, and *PIK3R6* (Supplementary Table S6; Fig 5). Both *NudE Neurodevelopment Protein 1 Like 1* (*NDEL1*) and *Phosphoinositide-3-kinase regulatory subunit 6* (*PIK3R6*) lie within the minimal 631 kb shared risk haplotype and contain variants demonstrating significant ASE in at least six of seven FCRs. *NDEL1* functions in neuron migration and neurite outgrowth, microtubule organization, and cell signaling. It has been associated with neurodegenerative disease (26) and may play a role in glioblastoma (27). *PIK3R6* functions in the PI3K/Akt pathway, which is commonly dysregulated in cancer, in leukocytes (28).

**Fig 5.**
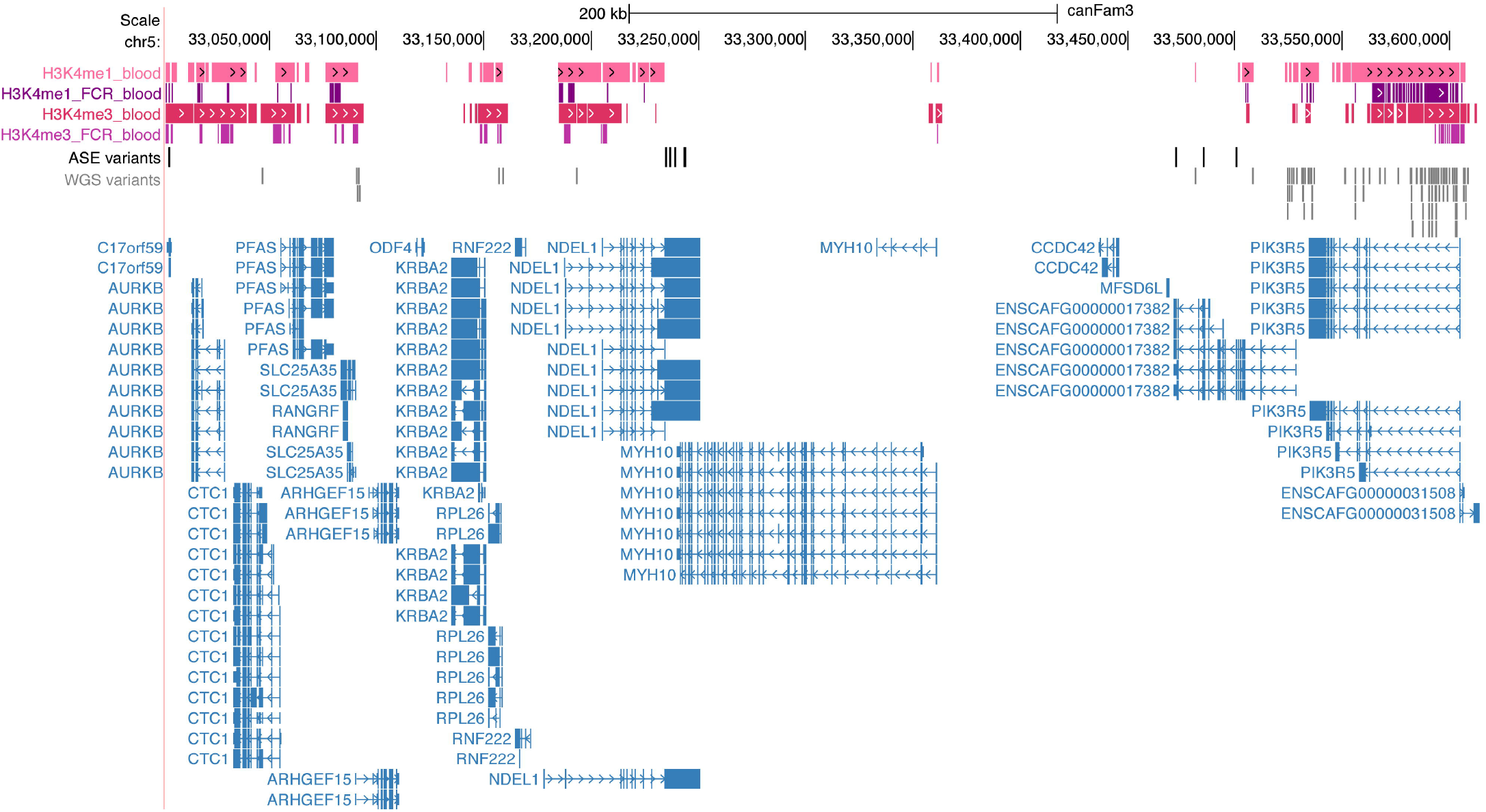
Multi-omics variant analysis. UCSC CanFam3.1 tracks at the 631 kb shared risk haplotype show the blood ChIP-seq regions for H3K4me1 and H3K4me3 for Bernese mountain dogs (pink) and the FCR (purple). ASE variants (black), 98 WGS variants meeting filtering criteria (gray), and the genes in the region are shown below (*ENSCAF00000017382* = *PIK3R6*).

As in human cancers, our data show histiocytic sarcoma is not fully explained by one gene or locus. We next tested effects of the CFA19 risk locus on changes in gene expression. We performed differential expression analysis between four cases homozygous for CFA19 risk and four heterozygous unaffected controls. The CFA5 genotypes were matched between the two groups with one dog homozygous for the CFA5 risk haplotype and three heterozygous dogs in each group. Among the top differentially expressed genes was *TNF alpha induced protein 6* (*TNFAIP6*; *P*_adjusted_=0.024), which lies 37 kb downstream of the GWAS susceptibility critical interval. *TNFAIP6* shows a 10.9-fold increased expression in histiocytic sarcoma cases homozygous for the CFA19 haplotype relative to heterozygous individuals. No other differentially expressed genes were proximal to the CFA19 critical interval (Supplementary Table S7). Three of the four cases demonstrate significant increased individual expression (z-score=2.84-7.61, equivalent to *P*<0.01) relative to all controls (n=7) at *TNFAIP6* (Supplementary Table S7). Comparison of the log2(TPM) expression at this gene for the four cases vs. seven controls indicates significant differential expression (Wilcoxon *P*=0.024); exclusion of the case who received chemotherapy increases the *P*-value to 0.067 (Supplementary Fig S2).

### Variant filtering and ChIP-seq analysis

We next sought to identify potential pathogenic variants within the CFA5 631 kb risk haplotype. Using WGS from four FCRs, three cases and one control, we filtered for variants concordant with the risk haplotype (See Table 2). Because FCRs and golden retrievers diagnosed with hematopoietic cancer shared a 631 kb risk haplotype, we hypothesized that they may also share the pathogenic variant(s) on this haplotype. We thus included published WGS from four golden retrievers diagnosed with B-cell lymphoma (three heterozygous for the risk haplotype and one homozygous) for filtering (Table 1). A total of 284 variants matched the segregation pattern of the CFA5 risk haplotype in the four FCR and four golden retriever WGS. A conservative allele frequency threshold of 50% in 1090 genomes from 233 other breed dogs (Supplementary Table S8) was applied to eliminate variants common across many breeds, resulting in 218 variants, none of which were unique to FCRs and golden retrievers (Supplementary Table S9). No variants were predicted to impact protein sequence or splice sites. Visual inspection of the interval in Integrative Genomics Viewer (29) revealed no structural variants segregating with the risk haplotype. The CanFam3.1 reference genome contains six gaps, totaling approximately 3 kb, within the critical interval, which may mask variants relevant to histiocytic sarcoma susceptibility.

We next investigated potential regulatory variants, which are not fully annotated in the CanFam3.1 reference. To identify promoter and enhancer regions in canine cell types relevant to cancers investigated herein, ChIP-seq data from peripheral blood mononuclear cells from seven dogs were generated for two histone marks, H3K4me1 and H3K4me3, to identify canine promoters and enhancers (See Methods, Supplementary Table S10). Publicly available ATAC-seq data identifying open chromatin regions from multiple canine tissues, i.e. spleen, lymph node, and bone marrow (30), were combined with blood ChIP-seq data to define regulatory regions.

Of the selected 218 variants (AF<50%) within the 631 kb critical interval, 98 overlapped with ChIP-seq and/or ATAC-seq regions (Fig 5). As none of the variants were completely unique to the FCR or golden retriever, we considered whether there could be combinations of variants private to the breed. The 98 variants were phased, allowing us to generate haplotypes in retriever and spaniel breeds, the latter of which were included because the retriever and spaniel clades share a recent common ancestor (16), yet the spaniel is not at risk for histiocytic sarcoma. Thus, a comparison of haplotypes in the region between the breeds might highlight combinations of variants that are neutral polymorphisms versus those that are unique to the retriever and possibly pathogenic. However, no blocks of continuous variants were unique to FCRs and golden retrievers. This does not preclude more distal combinations of variants that may be unique to affected individuals; however, it is likely that the causal mutations are present in other breeds.

### Transcription factor binding motif analysis and variant genotyping

To further explore candidate pathogenic variants, we selected regulatory regions from blood ChIP-seq data surrounding *NDEL1* and *PIK3R6*, candidates from ASE analysis, to interrogate variants for possible transcription factor binding motif alterations. ATAC-seq regions overlapped with blood ChIP-seq and were thus included. Regulatory elements at *PIK3R6* and *PIK3R5* contained 92% of the 98 variants overlapping ChIP-seq within the 631 kb shared haplotype (Fig 5, Supplementary Table S9). Five of the 98 WGS variants had significant scores in two transcription factor (TF) motif programs (See Methods), suggestive of a difference in binding affinity between the FCR risk and non-risk alleles (Supplementary Table S11); all were within *PIK3R5/6* regions. An additional variant (CFA5:33528647), significant in one TF binding affinity program (FIMO, *P*_adj_=0.0067) and demonstrating a difference in *SP1* and *KLF5* binding affinity between risk and non-risk variants in sTRAP (log(*P*)=0.5), was chosen for Sanger sequencing because it is within a human *PIK3R6* regulatory region in the GeneHancer promoter- and enhancer-gene interaction database where *SP1* and *KLF5* are reported to bind (31). Of the six variants selected for genotyping, one lies within a 12 bp G repeat in a GC-rich region, and we were unable to obtain reliable genotypes for this variant in all dogs (CFA5:33531804). The remaining five variants were significantly associated with histiocytic sarcoma (Supplementary Table S12). We calculated Fisher’s exact *P*-values for matched genotypes across 79 case and 69 control FCRs (Table 3). Variants at CFA5:33531780 and 33576022 had the lowest *P*-values of 4.2×10^−5^ and 4.6×10^−5^, respectively (lead SNP *P*=1.8×10^−4^, Table 3), and were located in ChIP-seq regulatory regions upstream of *PIK3R6*.

**Table 3.** Variants associated with histiocytic sarcoma and predicted to alter transcription factor binding sites.

| Position | Variant | *P*-value<sup>a</sup> | Odds Ratio | 95% Confidence Interval | AF in other breeds<sup>b</sup> | Transcription factor | Haplotype with preferred binding |
| --- | --- | --- | --- | --- | --- | --- | --- |
| 33528647 | C>CCG | $6.9 \times 10^{-5}$ | 2.63 | 1.64-4.21 | 0.29 | KLF5, SP1 | risk, risk |
| 33531780 | G>A | $4.2 \times 10^{-5}$ | 2.71 | 1.69-4.35 | 0.45 | SP1 | risk |
| 33576022 | GAGAA>G<br>GAAAGAA | $4.6 \times 10^{-5}$ | 2.64 | 1.65-4.23 | 0.011 | EWSR1-FL1 | risk |
| 33587141 | G>A | $1.2 \times 10^{-4}$ | 2.49 | 1.56-3.99 | 0.20 | IRF1 | risk |
| 33594214 | G>A | $1.1 \times 10^{-4}$ | 2.56 | 1.60-4.10 | 0.23 | PLAG1,<br>EWSR1-FL1,<br>RREB1 | non-risk,<br>risk, non-risk |

<sup>a</sup>Fisher's exact *P*-values in a matched population of 79 case and 69 control FCRs. Lead SNP in the same cohort $P=1.8 \times 10^{-4}$ , OR=2.48, 95%CI=1.55-3.97.
<sup>b</sup>WGS from 1066 breed dogs, excluding FCRs, golden retrievers, and Labrador retrievers.

The 631 kb shared haplotype delineated here is associated with histiocytic sarcoma in FCRs and B-cell lymphoma in golden retrievers, and we hypothesize that it harbors one or more pathogenic variants contributing to susceptibility for both diseases in each breed. Although less frequently than hemangiosarcoma and B-cell lymphoma, which affect 20% and 6% of the breed respectively (32), golden retrievers also develop histiocytic sarcoma (7% of all tumors in the breed) (4, 12, 33). Additional genotyping of the five candidate variants within this region, reveals that they are present in ~75% of FCRs with lymphoma (B-cell n=3, T-cell n=4, and unspecified subtype n=13, Table 1), and are in complete LD with the lead histiocytic sarcoma GWAS SNP (r^2^=1), consistent with our hypothesis. Three of the five variants were also in complete LD with this SNP in golden retrievers with B-cell lymphoma (n=9) or histiocytic sarcoma (n=21), i.e. CFA5:33576022, 33587141, and 33594214. The remaining two had r^2^ values of 0.32 (CFA5:33528647) and 0.16 (CFA5:33531780; Supplementary Table S12), indicating that they are not on the risk haplotype in golden retrievers. In aggregate, these data provide strong support that one or more of the three variants located on the CFA5 risk haplotype are likely to confer susceptibility to histiocytic sarcoma and B-cell lymphoma in both retriever breeds.

## Discussion

Genetic investigation of histiocytic sarcoma has been limited in humans due to its rarity, and identification of underlying genetic risk factors has not been undertaken. In this study, we leveraged the frequency of this cancer in the FCR breed and, in a GWAS of 309 dogs, identified two loci associated with histiocytic sarcoma on CFA5 and 19, respectively, that explain ~35% of risk. The former colocalizes with risk for two other cancers of hematopoietic origin and contains a shared risk haplotype for histiocytic sarcoma and B-cell lymphoma in FCRs and golden retrievers, respectively. We subsequently applied a multi-omics approach and identified ASE in *PIK3R6* and regulatory variants local to this gene that are predicted to impact TF binding. The second locus, on CFA19, has not been previously associated with cancer risk in dogs and increases risk for histiocytic sarcoma in combination with the CFA5 locus. The CFA19 critical interval lies upstream of *TNFAIP6* whose expression is upregulated in blood samples from FCR cases compared to healthy FCRs. This work reveals a shared germline predisposition for multiple hematopoietic cancers in dogs, reflecting clinical observations in the human literature of concurrent or subsequent histiocytic sarcoma with lymphomas thought to result from a common progenitor cell.

Chromosome 5 was recently identified, among other loci, through a histiocytic sarcoma GWAS in Bernese mountain dogs (34). In that GWAS, the best-associated SNPs were at 30.4 Mb. The authors noted that inclusion of 13 FCR cases and 15 FCR controls shifts their lead SNP to CFA5:33823740 with a slightly lower *P*-value, and they identify a common haplotype at 33.8-34.3 Mb (34). By comparison, our data strongly indicate that the primary risk haplotype for histiocytic sarcoma in FCRs is at 33.0-33.6 Mb and includes 18 genes. In our data, the maximum haplotype sharing is observed here among FCR cases (Fig 2C), as well as in the closely-related golden retriever breed, where cases diagnosed with B-cell lymphoma share the same haplotype (Fig 4). When genotypes for the CFA5 lead SNP were included as a covariate in our FCR histiocytic sarcoma GWAS, we also observed top SNPs proximal to CFA11:44 Mb and CFA2:29 Mb peaks (Supplementary Table S4) as described in the published Bernese mountain dog GWAS (34).

Our ASE RNA-seq analysis at CFA5 highlights two genes, the most compelling of which is *PIK3R6*, which encodes a regulatory subunit of the PI3Kγ heterodimer within the PI3K/Akt pathway. PI3Kγ expression is predominantly restricted to leukocytes (28) where it functions in cell migration, angiogenesis, and immune response (35). *PIK3R6* has a tumor suppressive role; knockdown increases PI3Kγ signaling and the potential for cells to metastasize (35, 36). The PI3K/Akt pathway is commonly activated in cancers (28), and somatic mutations in *PIK3CD* and *PI3KCA*, which encode other PI3K isoforms, have recently been detected in a subset of primary human histiocytic sarcoma tumors as well as secondary tumors and their co-occurring lymphomas (3, 37), although mutations in PI3Kγ genes have not yet been reported. *PIK3R6* was downregulated in B-cell lymphoma tumors from golden retrievers that harbored the CFA5:29 Mb hemangiosarcoma risk haplotype in this breed (19). Our ASE results suggest that variants on the CFA5:33 Mb haplotype may impact *PIK3R6* regulation.

Our ChIP-seq and WGS analyses identified five candidate variants strongly associated with histiocytic sarcoma in FCRs and predicted to impact TF binding sites proximal to *PIK3R6*. While these variants were also detected in FCRs and golden retrievers with lymphoma and golden retrievers with histiocytic sarcoma, supporting the hypothesis that variants within the shared risk haplotype underly susceptibility to both diseases in the retrievers, we note that sample sizes were small. Genotyping in larger cohorts is necessary to confirm the observations made here.

Among the candidate variants identified herein, CFA5:33576022, located upstream of *PIK3R6*, emerged as particularly strong, demonstrating the lowest allele frequency in 232 breeds that are at low to zero risk for developing histiocytic sarcoma (1%, Table 3). The variant consists of a GAA insertion within a GAAA microsatellite region and creates an ETS DNA binding site containing the core ETS 5’-GGAA-3’ motif. The ETS family of transcription factors are involved in tumorigenesis in many cancers, including lymphomas, leukemias, and Ewing sarcoma (38, 39). Functional studies will be necessary to elucidate the precise mechanisms by which the associated variants impact pathogenesis. We note there may be additional variants within the critical interval that contribute to risk through other mechanisms, such as post-translational modifications, or variants that function in a combinatorial fashion.

The CFA5 risk haplotype is sufficiently common among FCRs that the broader chromosomal region may have been under selection at some point. Multiple across-breed GWASs have associated the ~29-33 Mb region with skull shape (40-42), specifically muzzle length and breadth, both important factors in the FCR breed standard, i.e., ideal physical and behavioral breed characteristics (43). The deleterious cancer alleles may have increased in frequency in the FCR as a hitchhiking event with nearby variants that control the desirable skull shape phenotype.

At the second associated locus, CFA19, RNAseq analysis shows that *TNFAIP6* is upregulated in whole blood of cases homozygous for the CFA19 risk allele. This gene is immediately downstream of the critical interval defined by high LD. *TNFAIP6* is a member of the hyaluronan-binding protein family with roles in inflammation, extracellular matrix stability, and cell migration. Importantly, increased *TNFAIP6* expression is correlated with reduced survival for several cancers (44, 45), and it is upregulated in two lymphoma tumor types in humans (46). *TNFAIP6* is also a biomarker in colorectal cancer patients who have increased expression in peripheral blood cells relative to controls (47). Interestingly, the risk-associated allele at this locus was more common among cases in our study with periarticular tumors versus tumors at other sites at diagnosis.

At the CFA5 locus alone, similar proportions of cases and controls are heterozygous for the risk haplotype. But homozygosity for the CFA19 risk allele accounts for 88% of these cases (Table 2). The presence of some control dogs who are also homozygous for the CFA19 risk allele and heterozygous at CFA5 is likely due to the age cutoff of 11 years for enrolling controls (upper quartile of case age at diagnosis was 10 years, Supplementary Fig S1). While a higher minimum age for controls is ideal, most FCRs die by age 12 of cancer or cardiac, renal, or musculoskeletal disease (6). Late onset cancers have proven difficult to study in humans, e.g. prostate cancer. Our results demonstrate that it is possible to differentiate reliable case and control groups for late onset cancers in breed dogs, even though assignment to a control group will prove imperfect as the dogs age.

The most unexpected result in this study is the fact that together the CFA5 and 19 risk loci, explain ~35% of risk for histiocytic sarcoma in FCRs. This high value is due to the population structure of domestic dogs. Each breed is a closed population, experiencing frequent bottlenecks and strong artificial selection, resulting in small numbers of variants having large effects on traits. These results also highlight the value of the dog model for studies of complex cancers, particularly those with a heterogeneous phenotype. In the absence of family-based linkage studies, few mechanisms exist in human cancer genetics to identify risk loci for rare, but lethal, cancers.

Candidate genes at both loci, *PIK3R6* and *TNFAIP6*, have tumor suppressive and metastatic roles in other cancers, and mutations in members of the PI3K and tumor necrosis factor pathways have been identified in human histiocytic sarcoma and lymphoma tumors (3, 37). Our results suggest these pathways are also important in risk for developing these tumors, and that FCRs may be a valuable clinical model for the development of therapies for canine and human histiocytic sarcoma as well as other hematopoietic cancers.

## Materials & Methods

### Ethics Statement

Samples were collected with written informed owner consent in accordance with Animal Care and Use Committee guidelines at the collecting institution: National Human Genome Research Institute (NHGRI) Animal Care and Use Committee, GFS-05-1; Utrecht Animal Experiments Committee, as required under Dutch legislation, ID 2007.III.08.110; Colorado State University VTH Clinical Review Board, VCS #2019-227; and Departmental Ethics and Welfare Committee (University of Cambridge, protocol CR44).

### Sample Collection

DNA was isolated by standard phenol-chloroform protocol from whole blood samples of FCRs diagnosed with histiocytic sarcoma via histopathology or cytology, and FCRs age ≥10 years with no history of cancer (Supplementary Text S1). Samples originated from North America and Europe (Supplementary Table S13). DNA samples from FCRs diagnosed with lymphoma were included for variant genotyping, as well as golden retrievers with lymphoma or histiocytic sarcoma (Supplementary Table S13, Supplementary Text S1).

### Genome-wide association and haplotype definitions

Genotypes were generated on the Illumina (San Diego, CA, USA) Canine HD 170k SNP array for 177 FCR cases and 132 FCR controls (GEO GSE163784). Association analyses were performed with GEMMA (18). Illumina Canine HD 170k SNP array genotypes were downloaded for golden retrievers diagnosed with B-cell lymphoma (n=41) or hemangiosarcoma (n=143), and 172 who were ≥10 years old and cancer-free at collection (19). Additional details are in Supplementary Text S1. Risk haplotypes were defined from cases having at least one copy of associated alleles at lead GWAS SNPs. Centromeric and telomeric boundaries were delimited by SNPs at which ≥3 cases no longer shared at least one copy of the case-associated allele, and where this pattern extended beyond those boundaries for ≥1 kb. All genomic positions are reported in CanFam3.1.

### RNA-sequencing and expression analyses

RNA was isolated from 11 FCR whole blood samples (four histiocytic sarcoma cases and seven healthy, aged controls). For one of the four cases (FCR1), blood was drawn one week after beginning Lomustine chemotherapy treatment. The remaining three samples were collected prior to any treatment. Libraries were prepared using Illumina TruSeq Stranded Total RNA kit with Ribo-zero Globin depletion and sequenced to a minimum of 100 million pairs of 150 bp reads per sample on an Illumina NovaSeq6000 (SRA PRJNA685036). ASEReadCounter was used to determine allele counts for allele-specific expression analysis. Variants showing significant allele-specific expression (chi-square *P*≤0.05) in two or more individuals were prioritized. Read counts determined in RSEM were used for differential expression analyses in DESeq2 with Benjamini-Hochberg correction (48). Genes with adjusted *P*<0.05 and log2foldchange absolute values greater-than or equal-to 1 were considered significantly over- or under-expressed. To examine individual expression, z-scores were calculated at each gene by comparing the individual’s expression levels to the mean and standard deviation of the control group after variance stabilizing transformation of the counts in DESeq2 as described previously (22). Z-scores of +/- 2.5 indicate a significant change in expression (22). See Supplementary Text S1. All sequence data generated herein were aligned to the CanFam3.1 reference genome.

### ChIP sequence

Peripheral blood mononuclear cells were extracted by Ficoll (Cytiva, Marlborough, MA, USA) from fresh blood collected from two FCRs and six Bernese mountain dogs. As only two FCR samples were available for this experiment, we incorporated data from Bernese mountain dogs collected at the time of FCR sampling to identify general canine regulatory regions in cells of hematopoietic origin, which have not yet been fully annotated in the canine reference genome. Immunoprecipitation was performed for two histone marks, H3K4me1 and H3K4me3 (Supplementary Text S1; SRA PRJNA685036). ChIPseq regions identified in the FCR overlapped those in the Bernese mountain dog samples. Publicly-available ATAC-seq data from multiple breeds were also used to define regulatory regions from relevant tissues, including spleen, lymph nodes, and bone marrow (30). Megquier et al. (30) observed that patterns of enrichment for ATAC-seq sites were similar between individuals, regardless of breed, and different across tissues, as expected from studies in other species (49, 50).

### Whole genome sequence and variant filtering

Whole genome resequencing (WGS) data were generated (Supplementary Text S1), for five FCR histiocytic sarcoma cases and three healthy FCR controls ≥10 years old (SRA PRJNA448733 and PRJNA685036). Variants were filtered for concordance with associated risk haplotypes in FCRs and genomes of four golden retrievers diagnosed with lymphoma (19) (Supplementary Text S1). Allele frequencies were calculated from 1090 publicly available genomes from other breed dogs (Supplementary Table S8). Integrative Genomics Viewer (IGV), software for the visualization of genome sequence (29), was used to scan the CFA5 critical interval for large structural variants not called in the VCF file. Reads are color-coded in IGV to flag aberrant insert size indicating insertions, deletions, or interchromosomal rearrangements or pair-orientation indicating inversions, duplications, or translocations.

### Transcription factor motif analysis

For a given candidate variant, two fasta files were created, one containing the allele on the risk haplotype, the other the non-risk allele, each including 30bp on either side of the variant site. We required significant TF binding affinity predictions in two programs in order to increase confidence in the potential impact of a given WGS variant on TF binding. The first, Find Individual Motif Occurrences (FIMO) (51), scans input sequence for known transcription factor motifs, assigns a log-likelihood ratio score for each motif based on sequence position, converts these scores to P-values and q-values. Transcription Factor Binding Affinity Prediction (sTRAP) (52, 53) predicts binding affinity of each transcription factor in a given matrix to wild-type and mutant sequence, compares affinity values between the two, and calculates which TF has the greatest difference in affinity resulting from the sequence change. Variants for which the same motif was identified by both programs as significant in either the risk or non-risk allele sequence were prioritized (Supplementary Text S1).

### Variant genotyping

Genotyping was accomplished by Sanger sequencing or agarose gel electrophoresis. Primer sequences, thermal cycling, and reaction conditions are in Supplementary Table S14. PCR products were sequenced as described previously (54).

## Acknowledgements

We thank the NIH Intramural Sequencing Center for generating sequencing data, Andrew Hogan for DNA isolation and sample collection, Cathryn Mellersh and Jane Dobson for providing samples, David Sargan for critical reading of the manuscript, and the many owners and breeders who contributed samples and clinical data.

## Supplementary Information

**Supplementary Text S1**. Additional materials and methods.

**Supplementary Fig S1**. FCR GWAS cohort age distribution.

**Supplementary Fig S2**. Boxplots of transcripts per million counts for *TNFAIP6*.

**Supplementary Table S1**. FCR GWAS #1 results.

**Supplementary Table S2**. Haplotype analysis.

**Supplementary Table S3**. FCR GWAS #2 results.

**Supplementary Table S4**. FCR GWAS with CFA5 genotypes as a covariate.

**Supplementary Table S5**. Differential expression analysis results and individual expression z-scores for CFA5.

**Supplementary Table S6**. Allele-specific expression results for CFA5

**Supplementary Table S7**. Differential expression analysis results and individual expression z-scores for CFA19.

**Supplementary Table S8**. WGS SRA accession numbers.

**Supplementary Table S9**. WGS variant filtering.

**Supplementary Table S10**. Summary of ChIP-Seq read counts per sample.

**Supplementary Table S11**. Predicted effects of risk variants on transcription factor binding affinity.

**Supplementary Table S12**. Genotypes at candidate variants in retrievers.

**Supplementary Table S13**. Cohort information.

**Supplementary Table S14**. Primer sequences and reaction conditions for variant genotyping.

