## Supplemental Methods for "Multi-omics approach identifies germline regulatory variants associated with hematopoietic malignancies in retriever dog breeds"

*Sample Collection*

G.R. Rutteman and G.C.M. Grinwis performed pathology for all cases from The Netherlands. Morphological hallmarks of histiocytic sarcoma tumors include large round-oval or polygonal cells, sometimes spindle cells, and multinucleated giant cells, often with erythrophagocytosis (1). Overall, 86% of histiocytic sarcoma cases in the GWAS cohort were diagnosed by histopathology and 14% by cytology. While histopathology is ideal for diagnosis, cytology is reliable, with 84-96% of cytology cases confirmed as histiocytic sarcoma in a recent review of canine tumors (1). Sex was known for a subset of samples and was not significantly skewed between cases and controls (*P*>0.05).

FCRs used for variant genotyping (n=20) were diagnosed with a either B-cell, T-cell or unspecified lymphoma. Six Dutch FCRs were diagnosed by cytology with non-Hodgkin’s lymphoma, lacking a specific subtype identification; four were diagnosed with T-cell lymphoma by histopathology. Three American FCRs were diagnosed with B-cell lymphoma and seven with non-specified lymphoma; five had histopathology and 5 had unknown diagnosis methods. All cases received from CSU were diagnosed by histopathology: golden retrievers with B-cell lymphoma (n=8), multicentric lymphoma (n=1), or histiocytic sarcoma (n=6), and one flat-coated retriever with histiocytic sarcoma. One golden retriever from The Netherlands was diagnosed with histiocytic sarcoma by histopathology. Seventeen golden retrievers collected in North America were diagnosed with histiocytic sarcoma by a minimum of cytology.

*Genome-wide association and haplotype definitions*

After filtering for call rate ≥95% and minor allele frequency <0.01, 108 084 SNPs remained in the FCR (177 cases vs. 132 controls) SNP genotype dataset. GEMMA linear mixed model method was used for association analyses with standardized relatedness matrix to control for population substructure (2). In a GWAS using a subset of FCRs, 10 000 permutations were performed with a perl script to run GEMMA with random case-control assignments to establish significance thresholds. Principal component analysis was performed using eigensoft (3). Proportion of phenotype variance explained by genotypes was estimated with restricted maximum likelihood analysis in GCTA (4) with the estimated disease prevalence set at 17-20% (5) and likelihood ratio tests to calculate *P*-values.

Golden retriever SNPs were filtered as described for FCRs, leaving 115 119. One golden retriever with >5% missing genotypes was excluded. A total of 90 785 overlapping markers, after independent filtering in FCRs and golden retrievers, were used for haplotype analyses.

*RNAseq*

Whole blood (~2.5mLs) samples collected in PAXgene RNA tubes (PreAnalytiX, Hombrechtikon, Switzerland) were shipped to the lab within one day of the draw and stored at -80°C. RNA was isolated from thawed PAXgene tubes following the manufacturer’s protocols. RNA samples were quantitated and analyzed on an Agilent (Santa Clara, CA, USA) 2100 Bioanalyzer System or an Agilent 4150 TapeStation; all had RIN/RIN^e^ scores >7. Approximately 1 µg of RNA per sample was used for library preparation. RNAseq read quality was assessed in FastQC before and after preprocessing with Trimmomatic to remove adapter sequences and low quality bases (6). Trimmed reads were aligned to CanFam3.1 using the RSEM+STAR alignment protocol for RNA-seq quantification (7, 8) with the paired-end and strandedness reverse flags. Transcripts annotated with FEELNc were used for alignment (9).

*Allele-specific expression analysis*

VCF files were generated following GATK best practices. STAR (10) was used to align fastq files to the CanFam3.1 reference genome in two-pass mode. Read groups were added and duplicates marked with Picard tools with the READ_NAME_REGEX option set to null as recommended by GATK (11) for RNAseq data. Reads were split by exons and intronic overhang clipped with SplitNCigarReads. Variants from dbSNPv131 were used for base recalibration. HaplotypeCaller was used to call variants per sample, which were hard-filtered for FisherStrand (FS) > 30 and QualBy Depth (QD) < 2. Heterozygous sites with read depth ≥10 were included for ASEReadCounter. Reference or alternate counts <1% of the total read count or that were absent in genomes from 1090 breed dogs (Supplementary Table S2) were excluded from further analysis, as they were unlikely to be true heterozygous variants. A chi-square test (*P*≤0.05) was performed for each site in each individual.

*ChIPseq*

Fresh blood was drawn in yellow-top tubes with acid citrate dextrose anticoagulent at the owner’s veterinary clinic and shipped directly to the lab, where it was stored at 4ºC for a maximum of three days prior to FICOLL extraction. FICOLL®-paque plus procedures (Cytiva, Marlborough, MA, USA) were used to isolate peripheral blood mononuclear cells from whole blood samples. Cells were counted using a Cellometer Auto T4 (Nexelcom Bioscience, Lawrence, MA, USA). Two million cells were processed per sample. Chromatin immunoprecipitation was prepared using the SimpleChIP® Plus Enzymatic Chromatin IP kit (magnetic beads; Cell Signaling Technology, Danvers, MA, USA). We followed the protocol of the cell culture cross-linking and sample preparation, adjusting some parameters: 0.5 µl of micrococcal nuclease was used per IP, and a 250 µl volume of digested chromatin was sonicated on ice using a Vibra-cell^TM^ Ultrasonic sonicator VCX 130 (SONICS, Newtown, CT, USA) with a 1/8-inch probe and with the following settings: 20 sec ON/40 sec OFF/ 30% output for a total of nine minutes. Immunoprecipitation was performed using 2 µl of Abcam (Cambridge, United Kingdom) antibodies for H3K4me1 (rabbit polyclonal-AbCam, #ab8895), H3K4me3 (rabbit polyclonal-AbCam, #ab8580), and one negative control (IgG: Normal Rabbit IgG #2729) provided in the Cell Signaling kit. We also produced a 2% input sample. Quality controls were performed for each ChIP sample: 1) fragmented DNA was purified and 10 μl was separated by electrophoresis on a 1% agarose gel. The majority of chromatin was digested to one to five nucleosomes in length (150 to 900 bp). 2) Purified DNA was analyzed by quantitative real-time PCR using primer sets for PCR detection of the ribosomal protein L30 (RPL30) gene locus (F:5ʹ-TCTGAGGTAATGGAGGGAACC-3ʹ; R:5ʹ- GCAAGCGTTCGCTTCTAAA-3ʹ). Final DNA quantity was determined using Qubit quantification (ThermoFisher, Waltham, MA, USA) before sequencing. ChIPseq libraries were constructed from 10 ng of ChIP DNA using Ovation UltralowSystem V2 1-96 (TECAN, Männedorf, Switzerland) with 15 cycles of PCR amplification. The final libraries were purified twice using Agencourt AMPure XP PCR Purification Beads (Beckman Coulter, Brea, CA, USA). The libraries were pooled and then quantitated by qPCR. The pool balance was checked by performing a MiSeq run using a MiSeq Nano kit, version 2 (Illumina, San Diego, CA, USA). The percentage of each library in the pool was determined from demultiplexing and was used to rebalance the pool before sequencing on an Illumina NextSeq 550 using version 2 chemistry to achieve ≥28 million 75-base reads. Library preparation and sequencing were performed at the NIH Intramural Sequencing Center (NISC). The data were processed using RTA version 2.4.11. Reads were mapped to the Canfam3.1 genome reference using BWA-MEM (12, 13) and converted to BAM files using samtools 1.10 (14). ChIPseq peaks were quantified in each sample by counting the number of mapped reads overlapping the peak, using MACS2 2.2.1 (15). In order to create a track for each histone mark, we compared histone peaks between the six Bernese mountain dog samples using bedtools (intersect -c -sorted; merge) (16), keeping only peaks overlapping in at least two samples. Only one of the two FCR samples yielded enough ChIP DNA for sequencing. This sample was analyzed separately, thus we kept all peaks.

*Whole genome sequence and variant filtering*

Libraries were prepared with the Illumina TruSeq DNA PCR-Free Protocol (# FC-121-3001) and sequenced at NISC to ~40X coverage per DNA sample (150bp paired-end reads) on Illumina NovaSeq 6000. BAM and VCF files were generated as follows. Paired-end reads were aligned to CanFam 3.1 reference genome using the BWA-MEM algorithm (12), sorted with SAMtools (14), and screened for duplicate reads with PicardTools 2.9.2 (17). GATK 4.0.8.1 (18) was used for local realignment around novel indels based on published variants (19). Base quality recalibration in GATK was performed using dbSNP and Illumina Canine HD chip positions. Single nucleotide variants and small indels were called with GATK HaplotypeCaller in gVCF mode to call variants in individual genomes then jointly across genomes. Variants were filtered using bcftools (20). The VCF file included 1090 publicly available genomes from other breed dogs (Supplementary Table S8), which were used to calculate allele frequencies. Filtered variants were annotated using VEP/98 (20). Variants overlapping blood ChIP-seq peaks called from the six Bernese mountain dogs and the FCR and/or assay for transposase-accessible chromatin using sequencing (ATAC-seq) data from Megquier et al. (21) were identified with bedtools intersect (16).

*Transcription factor motif analysis*

JASPAR CORE vertebrates 2018 motifs and JASPAR vertebrates matrix, with human promoters background model, were used for FIMO and sTRAP, respectively. Significant binding affinity for either the risk or non-risk allele was defined in sTRAP as having p-value <0.05 and difference in affinity between the sequences of log(p)≥|0.5|. For FIMO, a Benjamini-Hochberg adjusted p-value ≤0.01 was considered significant where the TF was only called for the risk or non-risk allele sequence, but not both.
