## Supplemental Figure 1 for "Multi-omics approach identifies germline regulatory variants associated with hematopoietic malignancies in retriever dog breeds"

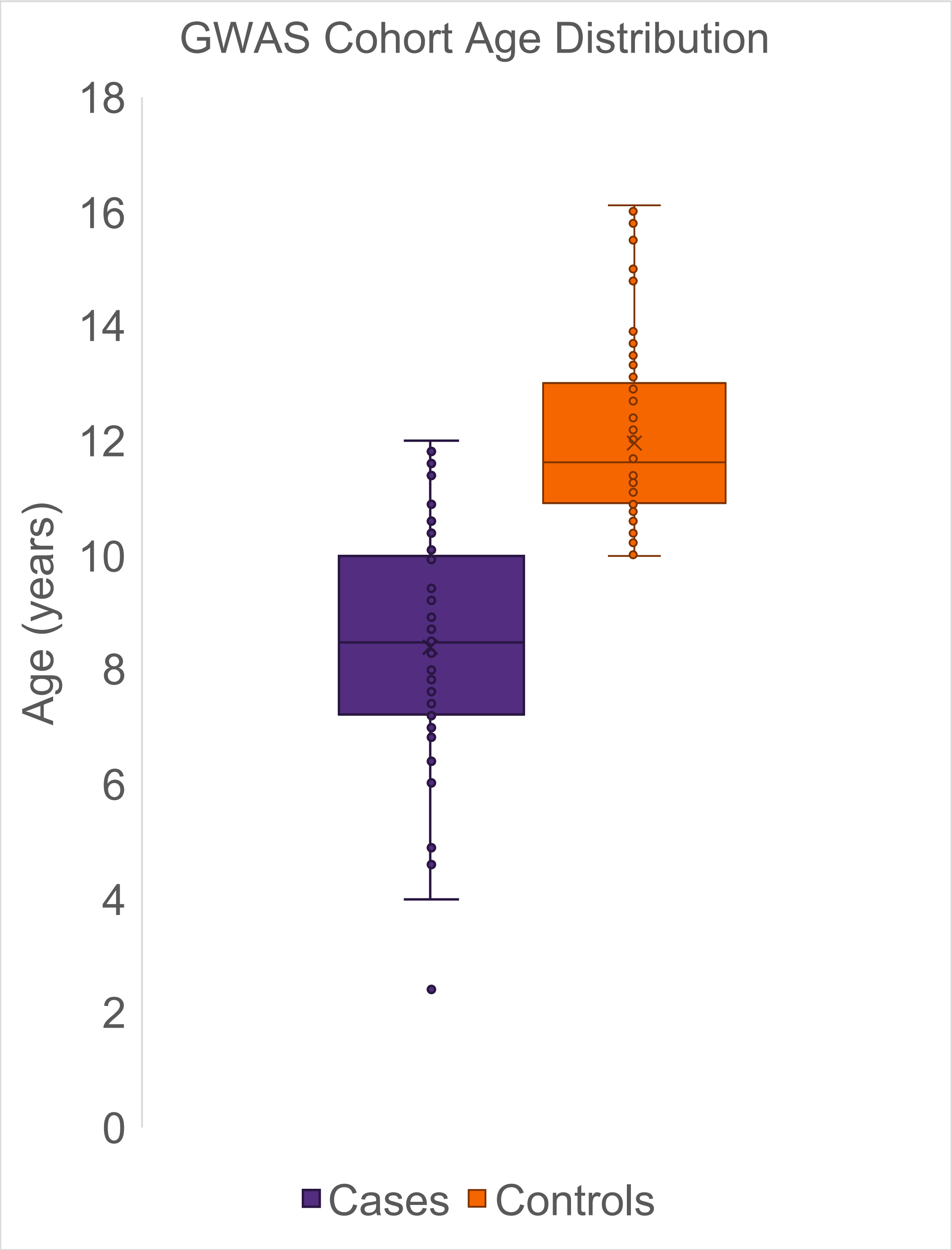


**Fig S1. Age of FCRs in the GWAS cohort.** Box plots are shown for case (n=68) age at diagnosis (left) and control (n=132) age at collection (right) with median ages of 8.5 and 11.6 years, respectively.
