## Supplemental Figure 2 for "Multi-omics approach identifies germline regulatory variants associated with hematopoietic malignancies in retriever dog breeds"

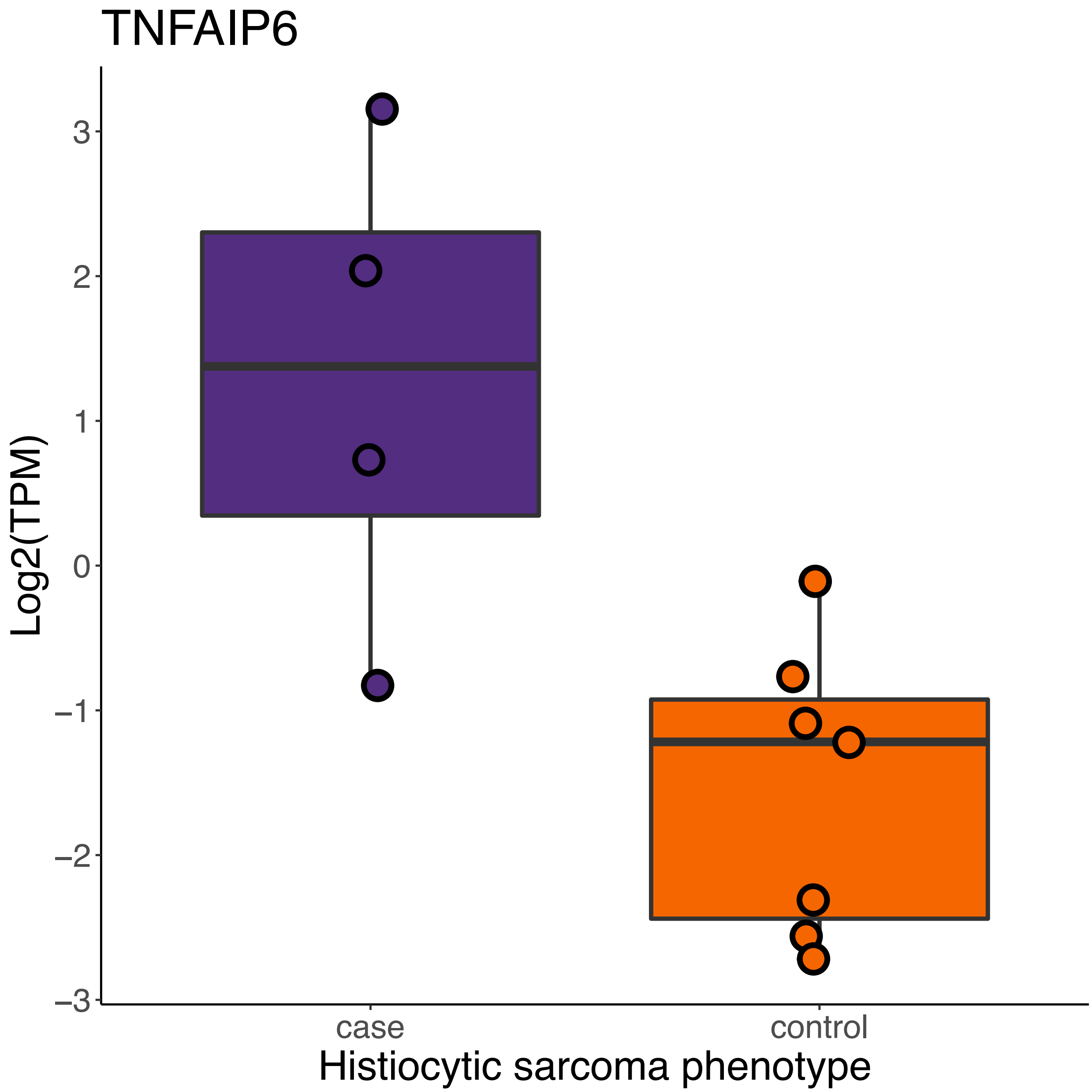


**Fig S2. Boxplots of transcripts per million counts for *TNFAIP6*.** Log2(transcripts per million) values are plotted on the y-axis for the four FCR cases homozygous for CFA19 risk (purple) and all seven FCR controls (orange), Wilcoxon *P*-value=0.024.
